# Exploring zebrafish larvae as a COVID-19 model: probable SARS-COV-2 replication in the swim bladder

**DOI:** 10.1101/2021.04.08.439059

**Authors:** Valerio Laghi, Veronica Rezelj, Laurent Boucontet, Maxence Frétaud, Bruno da Costa, Pierre Boudinot, Irene Salinas, Georges Lutfalla, Marco Vignuzzi, Jean-Pierre Levraud

## Abstract

Animal models are essential to understand COVID-19 pathophysiology and for pre-clinical assessment of drugs and other therapeutic or prophylactic interventions. We explored the small, cheap and transparent zebrafish larva as a potential host for SARS-CoV-2. Bath exposure, as well as microinjection in the coelom, pericardium, brain ventricle, bloodstream, or yolk, did not result in detectable SARS-CoV-2 replication in wild-type larvae. However, when the virus was inoculated in the swim bladder, a modest increase in viral RNA was observed after 24 hours, suggesting a successful infection in some animals. This was confirmed by immunohistochemistry, with cells positive for SARS-CoV-2 nucleoprotein observed in the swim bladder. Several variants of concern were also tested with no evidence of increased infectivity in our model. Low infectivity of SARS-CoV-2 in zebrafish larvae was not due to the host type I interferon response, as comparable viral loads were detected in type I interferon-deficient animals. Mosaic overexpression of human ACE2 was not sufficient to increase SARS-CoV-2 infectivity in zebrafish embryos or in fish cells in vitro. In conclusion, wild-type zebrafish larvae appear mostly non-permissive to SARS-CoV-2, except in the swim bladder, an aerial organ sharing similarities with the mammalian lung.

## Introduction

The COVID-19 pandemic has taken an enormous toll worldwide, both in human and economic losses. Although vaccination is finally under way, the SARS-CoV-2 virus is predicted to persist for years (Moore et al., 2021), and its variants represent an unpredictable threat. Thus, it will be necessary to continue the research efforts to understand its heterogeneous pathology and develop new drugs and vaccines.

Animal models play a central role during any pandemic since they are essential to analyze pathology, transmission, and test vaccines and drugs. Besides non-human primates, other mammals such as Syrian hamster and ferrets are naturally susceptible to SARS-CoV-2 (Muñoz-Fontela et al., 2020). Mice, the most widely used model for host-pathogen studies, require transgene-mediated expression of human angiotensin converting enzyme 2 (hACE2) to be infected (Lutz et al., 2020), although some recent variants replicate to a significant extent in wild-type mice (Montagutelli et al., 2021). All these models have several advantages and disadvantages. Non-human primates are very expensive, require large animal facilities and are not conducive to large scale experiments. hACE2 transgenic mice remain expensive and not readily available. As a result, expanding the repertoire of animal models for any disease is always beneficial and each model may shed light to unique aspects of the pathogen-host interaction. Here, we test if zebrafish larvae can be added to the list of suitable animal models for the study of COVID-19.

The zebrafish larva is an increasingly popular model to understand host-pathogen interactions (Torraca & Mostowy, 2018). Low cost of husbandry, high fecundity, and small size and transparency at early stages are among its main advantages. Thus, zebrafish larvae allow live imaging of pathogen dissemination at the whole organism to subcellular scales, and *in vivo* molecule screens in 96 well formats. Zebrafish is also a genetically tractable model, and thousands of mutant and reporter transgenic lines are available in fish facilities and repositories worldwide. Given that 80% of disease-associated genes of humans have a zebrafish orthologue (Howe et al., 2013), it is not surprising that zebrafish continue to be developed as models for human pathogens. Further, zebrafish is a bony vertebrate with an immune system that is also highly similar to ours. For instance, orthologs of the classical inflammatory cytokines (IL1β, TNFα, IL-6) as well as type I interferons (IFNs) are all found in zebrafish (Zou & Secombes, 2016). Interestingly, zebrafish adaptive immunity develops only at the juvenile stage, weeks after hatching (Lam et al., 2004), and the larva thus constitutes a system where innate immunity can be evaluated independently of adaptive responses. These assets make the zebrafish highly suitable to the study of host-virus interactions (Levraud et al., 2014).

Experimental infection has been established with various human viruses in zebrafish, including Herpes simplex virus 1 (Burgos et al., 2008), Chikungunya virus (CHIKV) (Palha et al., 2013), Influenza A virus (IAV) (Gabor et al., 2014) and norovirus (Van Dycke et al., 2019). The upper temperature limit of proper zebrafish development, 33°C (Kimmel et al., 1995), may be an issue for some viruses; however, it corresponds to that of upper airways, and in fact SARS-CoV-2 replicates better at 33°C than at 37°C (V’kovski et al., 2021). The absence of lungs is another drawback to model a respiratory infection; however, teleost fish do possess an air-filled organ, the swim bladder, used for buoyancy regulation. Lungs of tetrapods and swim bladders of fish are evolutionary related and share important structural homologies, such as surfactant coating (Cass et al., 2013). In support, inoculation of IAV in swim bladder resulted in localized infection (Gabor et al., 2014).

The zebrafish genome contains a unique, unambiguous ortholog of the gene encoding ACE2, the SARS-CoV-2 receptor; however, modest conservation of amino-acids at the binding interface make fish ACE2 proteins unlikely to bind the virus spike efficiently (Damas et al., 2020). Despite these *in silico* predictions, host susceptibility requires experimental validations, especially given that many other receptors and co-receptors for SARS-CoV-2 have been identified (Zamorano Cuervo & Grandvaux, 2020). In zebrafish larvae, based on single cell transcriptomics, *ace2* is strongly expressed in a subtype of enterocytes (Postlethwait et al., 2021); the gut is also the organ with strongest *ace2* expression in humans.

There have been reports of the use of zebrafish to study COVID-19. We have recently reported pathological effects after exposure of zebrafish to recombinant SARS-CoV-2 spike protein, including accelerated heart beat in larvae and severe olfactory damage causing transient hyposmia in adults after intranasal administration (Kraus et al., 2020). Injection of recombinant spike to adults has also been reported to induce adverse effects (Ventura Fernandes et al., 2020). Xenotransplantation of human lung cells in the swim bladder of adult zebrafish has been proposed to test the effect of an herbal drug on SARS-CoV-2 (Balkrishna et al., 2020). However, to date, no in-depth assessment of the ability of SARS-CoV-2 to replicate in zebrafish has been published.

Here we tested several tactics to infect zebrafish larvae with SARS-CoV-2, including bath exposure and microinjection in various organs or cavities. The swim bladder was the only organ that supported SARS-CoV-2 replication in wild-type larvae. Preventing type I IFN responses did not result in increased replication, consistent with the fact that SARS-CoV2 inoculation did not result in strong IFN responses or induction of inflammatory cytokines.

## Results

### SARS-CoV2 replicates in zebrafish larvae only when injected in the swim bladder

We first tested if an early strain of SARS-CoV-2 would replicate in wild-type zebrafish larvae after bath exposure. We exposed 4 days post fertilization (dpf) larvae with inflated swim bladders (ensuring an open gut) as well as 2 dpf dechorionated embryos with suspension of either live or heat-inactivated virus added to water (8×10^4^ PFU/mL). Larvae were then incubated at 32°C and observed regularly; no specific signs of distress were noted. After RNA extraction, the amount of polyadenylated SARS-CoV2 *N* transcripts were measured by qRT-PCR. Although viral RNA was readily detectable after 6 hours of exposure, it then declined and became undetectable after 48 hours (Figure 1). Therefore, bath exposure failed to achieve infection.

**Figure 1.**
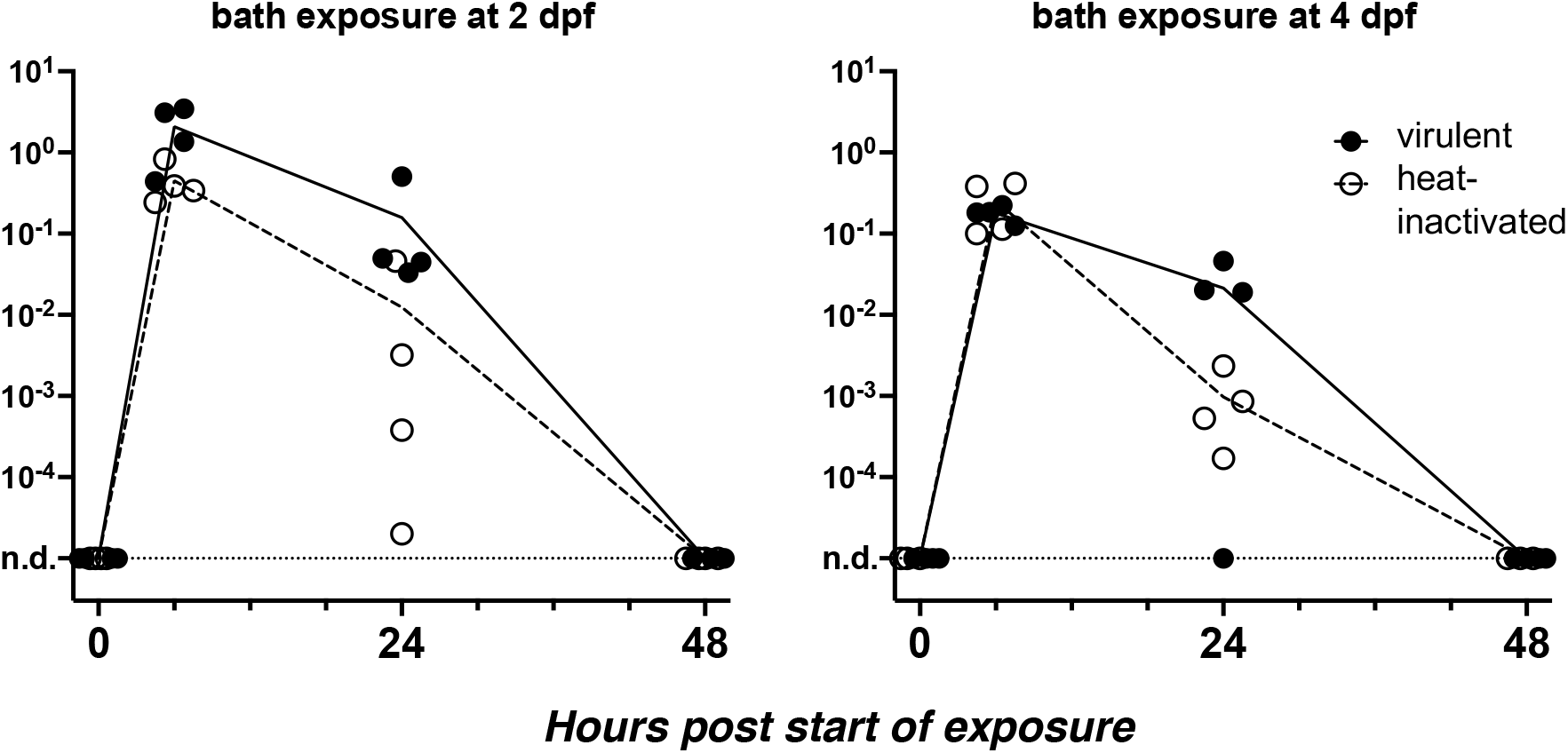
Bath exposure of zebrafish larvae to SARS-CoV2. Kinetics of qRT-PCR measurements of polyadenylated viral N copies; each point corresponds to an individual larva. N.d., not detected.

We then turned to microinjection of larvae with SARS-CoV-2. Using a camera-fitted macroscope under a biosafety hood, a concentrated SARS-CoV-2 suspension was microinjected in various sites of 3 dpf larvae (Figure 2A). Compared with our previous experience of microinjection using the eyepieces of a stereomicroscope, this was significantly harder, notably due to lack of stereovision. These challenging injection conditions resulted in variability during early attempts; this later improved greatly, and although success of intravenous (IV) injections remained difficult to ascertain, others, notably in the coelomic cavity, were achieved reliably and in a reasonable time frame. Injection of the syncytial yolk cell was relatively easy, but leakage was often observed after capillary withdrawal, in which case larvae were discarded. Injected larvae were immediately rinsed and transferred into individual wells of 24-well plates, which were then incubated at 32°C. Larvae were imaged daily; none of the typical disease signs that we noted during other viral infections (e.g., edemas, spine bending, necrotic spots, slow blood flow) (Palha et al., 2013) were observed.

**Figure 2.**
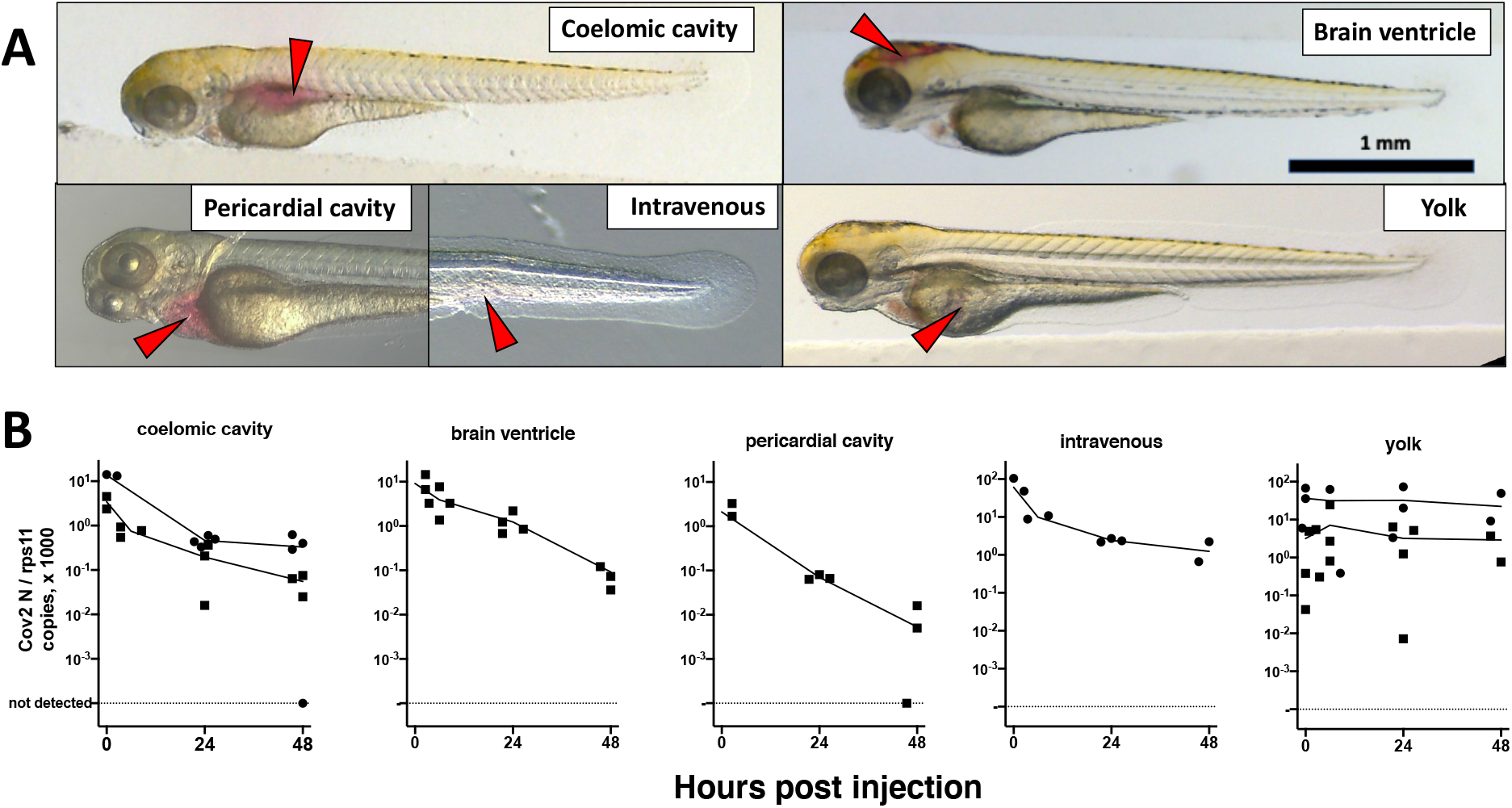
Microinjection of SARS-CoV2 to 3dpf wild-type larvae. A. Illustrations of the targeted sites. Images taken less than one minute after injection of the phenol red-coloured SARS-CoV-2 suspension. Red arrowheads point to the sites of microinjection. B. quantification of polyadenylated N transcripts over time, assessed by qRT-PCR; each symbol is an individual larva. Circles and squares correspond to injection of viral suspensions 1 and 2, as labelled on Table 1, respectively. Lines connect the means of values measured at each time point.

At various time points, individual larvae were euthanized and RNA extracted. The initial inoculum, measured in larvae lysed ∼30 minutes post-injection (pi), was readily detectable by qRT-PCR (Table 1). Absolute quantification by qRT-PCR, using certified commercial reagents, revealed an amount of polyadenylated SARS-CoV-2 *N* transcripts that was ∼10^4^-fold higher than the injected number of PFU (Table 1). Therefore, the overwhelming bulk of viral RNA injected in larvae must correspond to non-infectious molecules.

**Table 1.** initial sense N copy numbers. Quantification by RT-qPCR of polyadenylated viral N transcripts in zebrafish larvae microinjected with 2nL of viral suspension (diluted 1.1-fold by addition of phenol red) in the coelomic cavity less than one hour before lysis.

|  | Viral suspension 1 | Viral suspension 2 |
| --- | --- | --- |
| Titer (PFU/mL) | 1.13 x 10<sup>8</sup> | 1.6 x 10<sup>7</sup> |
| PFU in 2nL inoculum | 205 | 29 |
| Median *N* copies measured in a cDNA sample corresponding to 1/100<sup>th</sup> of larval extract | 11026 | 5679 |
| 95% confidence interval | 5175-12255 | 4967-7854 |
| Number of samples | 23 | 12 |
| Ratio of median *N* copies to PFU | 5378 | 19583 |

We then measured polyadenylated *N* copies over time. A decline was observed for all injection sites, with the notable exception of the yolk (Figure 2B). Amounts measured in yolk were highly variable at early time points, more than in other sites, probably due to leakage.

To determine if the relatively high amounts detected in yolk at late time points were due to viral replication, we re-analyzed these RNA samples by performing reverse transcription with a primer that hybridizes to the 5’ leader sequence of negative strand subgenomic RNAs, a hallmark of active SARS-CoV-2 replication (Kim et al., 2020; Wölfel et al., 2020). Such transcripts were detected in the initial inoculum, but in lower amounts than polyadenylated transcripts (median values of 1042 and 191 copies for coelom-injected larvae with viral suspensions 1 and 2, respectively). In coelom-injected larvae, these antisense transcripts decreased and became undetectable at 48 hours post-injection (hpi). By contrast, in yolk-injected larvae, levels were stable (Figure S1A). Therefore, both sense and antisense viral RNA molecules appeared to be protected from degradation in the yolk, and there was no clear evidence for viral replication. Notably, we did not observe yolk opacity in injected animals, a hallmark of yolk cell infection with other viruses such as CHIKV (Palha et al., 2013) and Sindbis virus (SINV) (Figure S2).

We then tested microinjection of SARS-CoV-2 in the swim bladder, which inflates at 3.5-4dpf (Parichy et al., 2009). We noticed that when the liquid was injected at the rostral end of the bladder, it was rapidly expelled via the pneumatic duct connecting the swim bladder to the esophagus. By contrast, when liquid accumulated at the caudal end of the swim bladder, if was well retained (Figure 3A). Therefore, injections were performed at 4dpf by targeting the caudal half of the bladder; larvae with liquid injected at the rostral pole were discarded. As age-matched controls, we also injected 4dpf larvae in the coelomic cavity, *i*.*e*. just next to, but outside of the swim bladder (Figure 3A)

**Figure 3.**
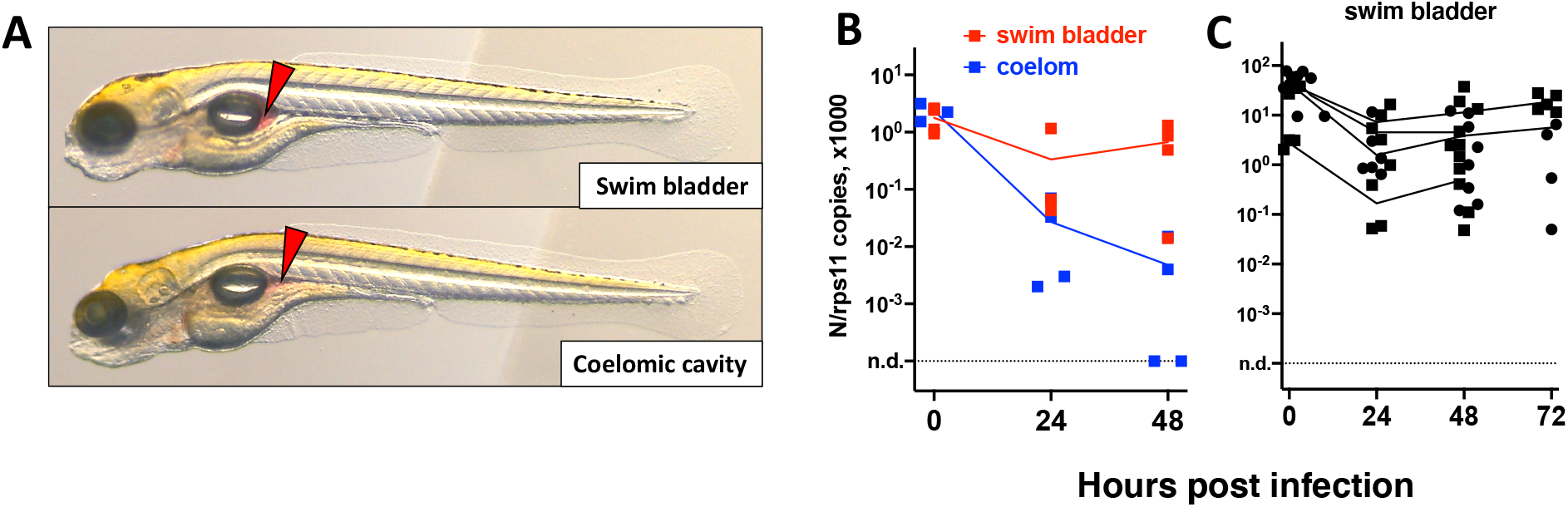
Microinjection of SARS-CoV-2 to 4dpf larvae. A. illustrations of injection in the posterior end of the swim bladder or in the coelomic cavity. B-C. quantification of polyadenylated N transcripts over time, assessed by qRT-PCR; each symbol is an individual larva. B. comparison of swim bladder (red) and coelom (blue) injection in a single experiment. C, four more swim bladder injection experiments. Lines connect the means of values measured at each time point Circles and squares correspond to injection of viral suspensions 1 and 2, as labelled on Table 1.

Remarkably, after an initial decrease of viral transcripts during the first 24 hours, a subsequent increase was often noted in swim bladder-injected larvae; by contrast, the decline continued in coelom-injected larvae (Figure 3B). This suggests that in swim bladder, after an initial degradation of viral transcripts, *de novo* production is taking place, implying successful infection. However, no disease signs were observed. We repeated the swim bladder injection several times finding consistent results; extending the experiment by one day yielded comparable results at days 2 and 3 (Figure 3C). We also measured antisense transcripts in these larvae, observing the same trend (Figure S1B).

To perform statistical analysis with reasonable power, we normalized the results of each independent experiment to the mean of the values measured just after inoculation, and then pooled the results by injection type. Because the dispersion increased considerably with time, we performed tests that allowed for unequal SDs when comparing time points. This analysis confirmed that after injection in the coelomic cavity, viral RNA amounts decline from 0 to 24 hpi and again from 24 to 48 hpi. By contrast, values measured in yolk were stable. In the swim bladder, a very significant decrease is observed during the first 24 hpi; while from 24 and 48 hpi, a non-significant re-increase of the means is observed (Figure 4A). Comparison between the coelom and the swim bladder showed a significantly higher level of viral RNA in the latter at 48 (but not 24) hpi (Figure 4B), consistent with a successful infection in the swim bladder.

**Figure 4.**
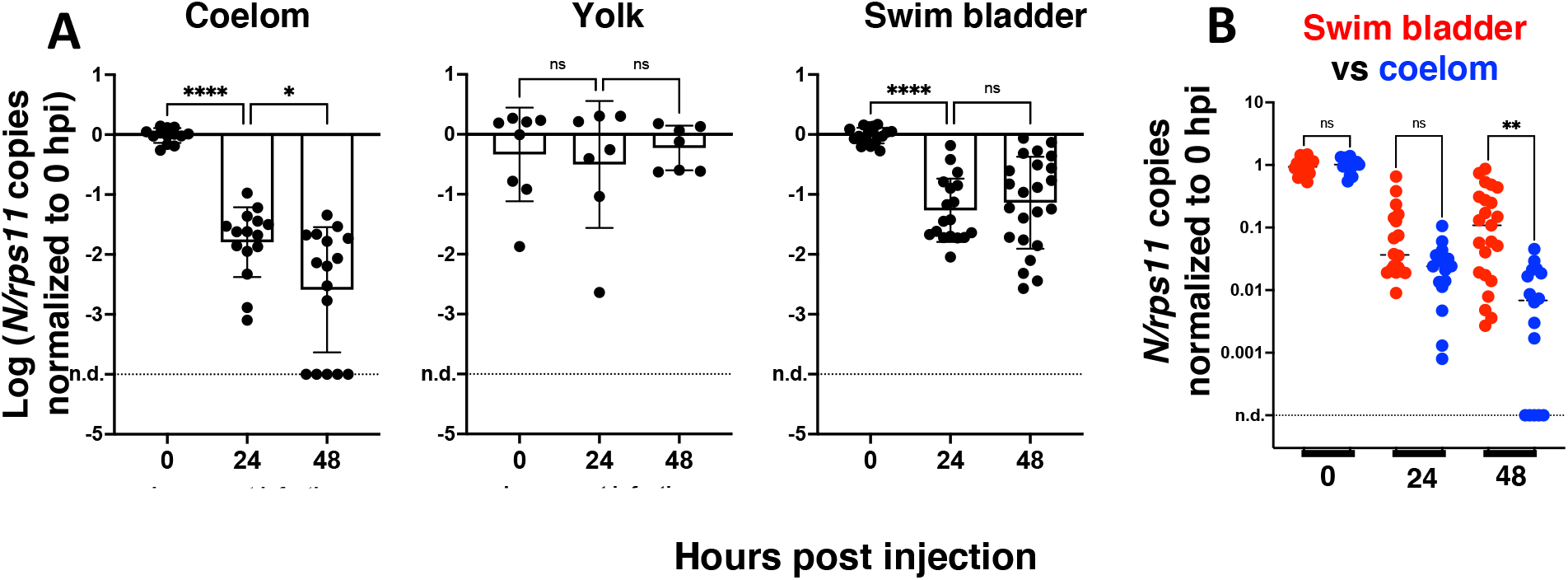
Statistical analysis of viral transcript quantifications. A. Comparison of SARS-CoV-2 RNA loads over time in each microinjection location; ANOVA analysis of log-transformed values, not assuming equal SDs (Brown-Forsythe test with Dunn’s correction). B. Comparison of coelom and swim bladder injections at each time point; non-parametric multiple comparisons of non-transformed values (Kruskal-Wallis test with Dunn’s correction). Ns, not significant; *, p<0.05; **, p<0.01; ****, p<0.0001. Results pooled from four, two and five experiments for coelom, yolk and swim bladder injections, respectively, after normalization to the means of values measured at 0 hpi for each experiment.

To confirm infection by SARS-CoV-2 after SB injection, we used whole-mount immunohistochemistry (WIHC). We tested several commercial Abs against the SARS-CoV-2 nucleoprotein, and selected a mouse Mab with minimal non-specific staining of naïve larvae, except for dots in the notochord that we routinely observe and are due to the secondary antibody only (Levraud et al., 2009). As an anatomical reference, we also labelled glial fibrillary acidic protein (GFAP), to reveal glial cells and main nerves. In most virus-inoculated larvae at 2 dpi, a patchy signal for N could be clearly detected in the swim bladders which were partially collapsed due to the fixation and staining procedure (Figure 5). 3D reconstruction (movie S1) indicate that these signals correspond to a few infected cells in the bladder wall, generally located close to the rear pole. No infected cells were detected outside of the swim bladders.

**Figure 5.**
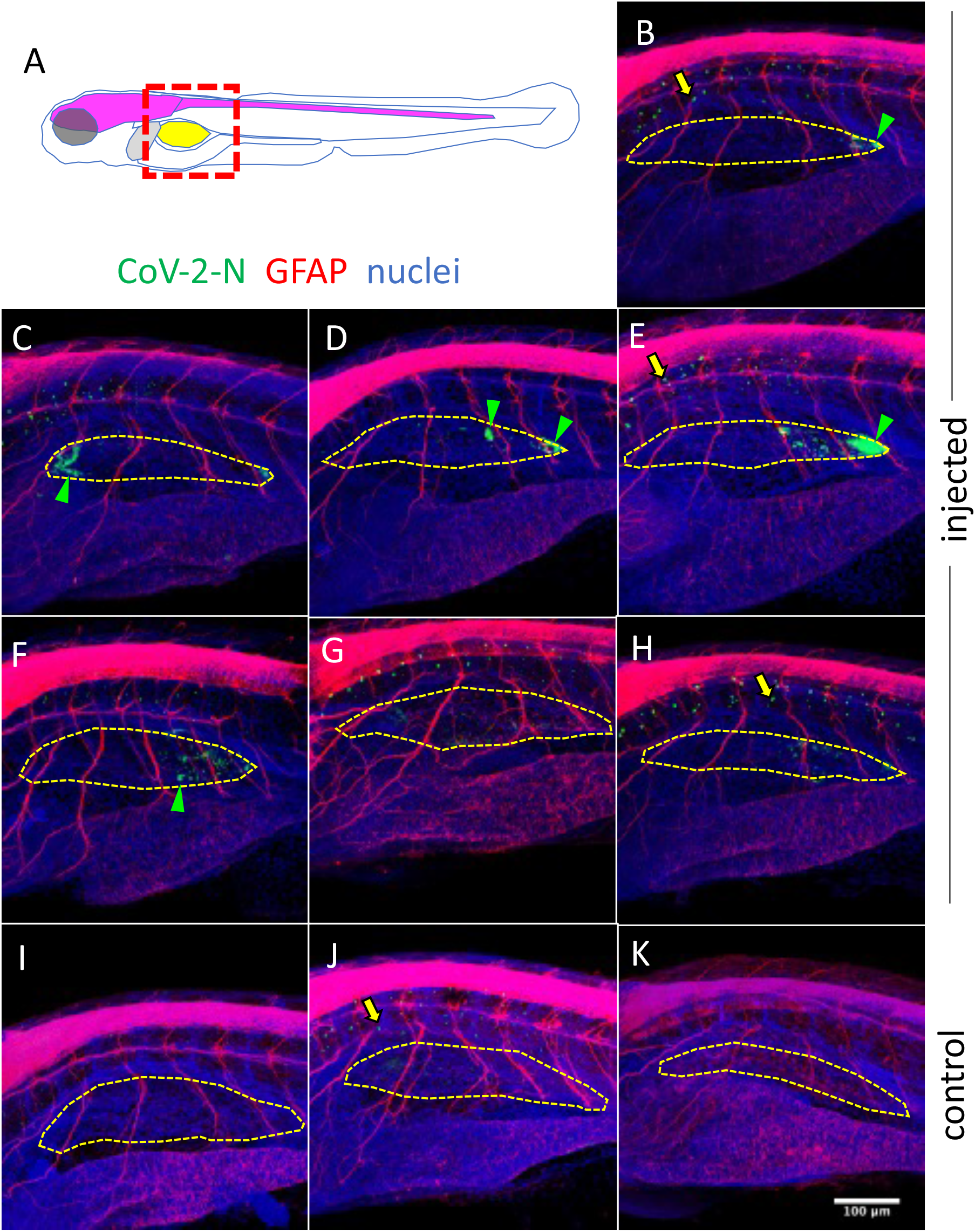
Immunodetection of infected cells in the swim bladder. A. scheme of the imaged region: the swim bladder is shown in yellow, the brain and spinal cord in magenta, the liver in grey. B-K, confocal images of SARS-CoV-2 injected (B-H) or uninjected (I-K) larvae fixed at 2 dpi and subjected to whole mount immunohistochemistry with an anti CoV-2-N antibody (green) and an anti-GFAP antibody (red), and with nuclei shown in blue. Maximal projections. The approximate contours of the partially collapsed swim bladders are shown with a dotted yellow line. N-positive cells shown with green arrowheads. Yellow arrows point to non-specific punctate signal in the notochord.

### Variants of concern do not show increased infectivity in wild type larvae

We then tested a series of SARS-CoV-2 variants by swim bladder inoculation. We obtained aliquotes from early passages after isolation of clinical strains, which had been titered at 3.10^7^ PFU/mL or more and thus did not require further concentration. We tested the alpha variant (formerly known as UK variant, or B1.1.7), the beta variant (South African variant, B1.351), the gamma variant (Brazilian variant, P1) as well as a representative of the G-clade which arose early during the pandemic. Non-diluted viral suspensions were injected as described above in the swim bladder of 4dpf larvae, and were then monitored for two days; no clinical signs were observed. Viral replication was assessed by qRT-PCR. A global decline of polyadenylated N transcripts over time was observed with all variants (Figure 6). One unique larva injected with the gamma variant was found to contain slightly more N copies than the initial inoculum; therefore, the experiment was repeated for the gamma variant, and again, one larva did not show the same decline as others. Thus, results obtained with the gamma variant were comparable to those obtained with the initial strain, with a fraction of larvae in which some replication appeared to take place. No replication was found with the other strains, which also corresponded to lower inocula according to qPCR results. Overall, we saw no evidence for an increased infectivity of SARS-CoV-2 variants in zebrafish larvae.

**Figure 6.**
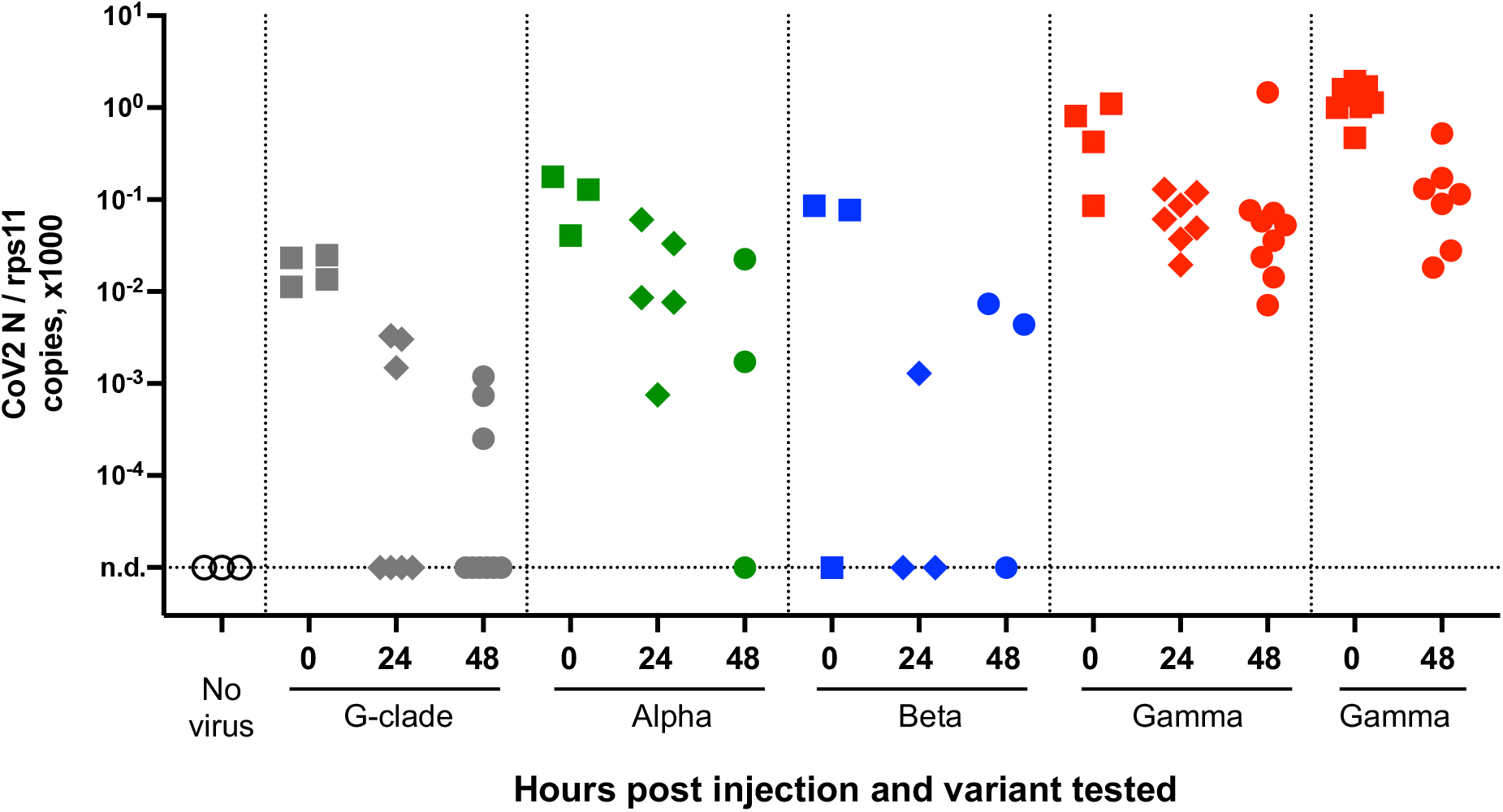
Testing SARS-CoV-2 variants. qRT-PCR analysis of larvae at various times after injection of 2nL of virus suspension in the swim bladder. Dotted lines separate independent experiments.

### Hours post injection and variant tested

### A defective type I interferon response does not increase SARS-CoV-2 replication

Type I interferons (IFNs) are key antiviral cytokines in vertebrates, including teleost fish. We thus tested if SARS-CoV-2 may replicate in larvae with a crippled type I IFN response.

First, we used morpholino-mediated knockdown of the type I IFN receptor chains CRFB1 and CRFB2, known to make zebrafish larvae hypersusceptible to infection with CHIKV or SINV (Boucontet et al., 2018; Palha et al., 2013). After injection of SARS-CoV-2 in the coelom of 3dpf larvae, decline of N transcripts was found to be similar in IFNR-knocked down larvae than in controls (Figure 7A).

**Figure 7.**
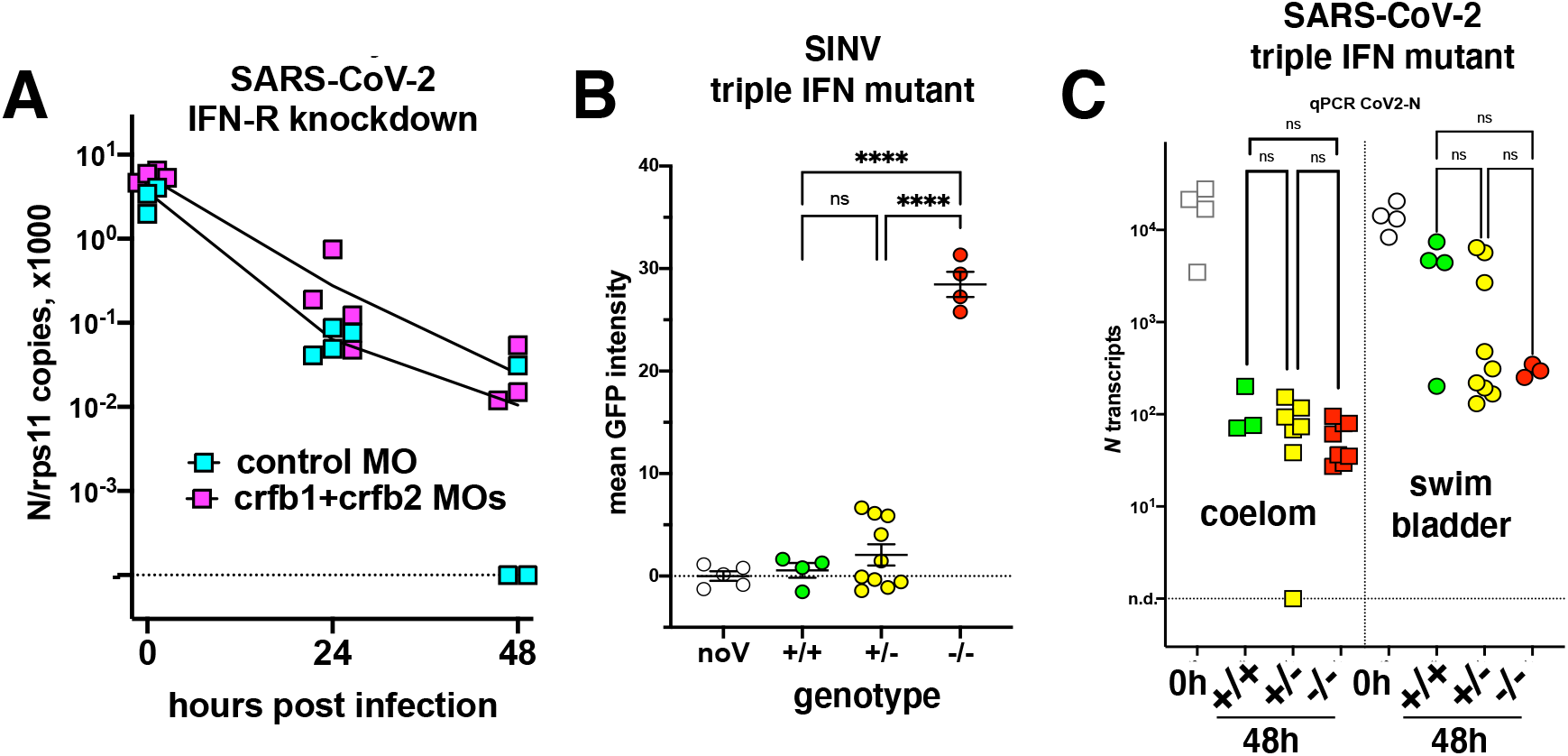
viral infection in IFN-defective larvae. A. IFN-receptor (crfb1 and crfb2 genes) or control morphants infected at 3dpf in the coelomic cavity; qRT-PCR. B and C. offspring from an incross of heterozygous triple IFN-mutants. B larvae injected with SINV-GFP IV at 3dpf, analyzed by fluorescence imaging at 48hpi. C. larvae injected with SARS-CoV-2, either at 3dpf in the coelom, or at 4dpf in the swim bladder; analysed by qRT-PCR at 0 or 48hpi. B and C, analysis by 1-way ANOVA.

To ensure a long-lasting suppression of the IFN response, we used a newly generated mutant zebrafish line dubbed “triple ϕ”, in which the three type I IFN genes *ifnphi1, ifnphi*2, and *ifnphi*3, tandemly located on chromosome 3, have been inactivated by CRISPR. Heterozygous triple ϕ mutants were viable and fertile; incrossing them yielded homozygous embryos at the expected mendelian ratio of ∼25%. Homozygous triple ϕ mutants could be raised up to juvenile stage, but, unlike their siblings, died in the two weeks following genotyping by fin clipping. To validate the phenotype of the mutants, we injected SINV-GFP to 3 dpf larvae from a heterozygous incross. 48 h later, all larvae were alive although some showed strong signs of disease, including loss of reaction to touch, abnormal heart beating, slow blood flow, edemas and opacified yolk spots. All larvae were imaged with a fluorescence microscope to measure the extent of infection, then lysed individually and genotyped. Homozygous mutant displayed a considerably higher level of fluorescence (Figure 7B), and were also identified *a posteriori* as the sickest larvae, confirming that triple ϕ mutants are hypersusceptible to viral infection.

Larvae from triple ϕ heterozygous incrosses were thus injected with SARS-CoV-2, either in the coelomic cavity at 3dpf or in the swim bladder at 4dpf. Larvae were lysed at 48hpi, analysed by qRT-PCR, and genotyped. Consistent with previous results, a 100-fold decrease of viral RNA was observed in coelom-injected larvae, while a weaker decrease was observed for swim bladder injection, with a bimodal distribution suggesting that infection happened in about one third of cases. In both situations, viral loads in homozygous triple ϕ mutants were not different from their wildtype siblings (Figure 7C). Thus, our results indicate that type I IFN responses are not responsible for the lack of replication of SARS-CoV-2 observed in wild-type zebrafish larvae.

### Lack of detectable inflammatory responses in SARS-CoV-2 injected larvae

We then tested if SARS-CoV2 inoculation in the swim bladder resulted in induction of a type I interferon response or inflammatory cytokines. For this, we performed qPCR on dT17-primed cDNAs from whole larvae. Based on our previous results (Kraus et al., 2020; Levraud et al., 2019), we tested the main type I interferon genes inducible in larvae, namely *ifnphi1* and *ifnphi3*; the strongly IFN-inducible gene *MXA*; the classical inflammatory cytokines *il1b* and *tnfa*; cytokines that reflect induction of type 2 or type 3 responses, *il4* and *il17a/f3*, respectively, and chemokines *ccl19a*.*1* and *ccl20a*.*3*. Although individual experiments suggested significant induction of *ifnphi1* at 48hpi or *il17a/f3* at 72h, this could not be replicated; as shown on Figure 8, in which data from 4 independent experiments have been pooled, no significant change in expression of any of these genes can be observed compared to uninjected control larvae. Similar negative results were obtained with larvae injected at different sites (not shown).

**Figure 8.**
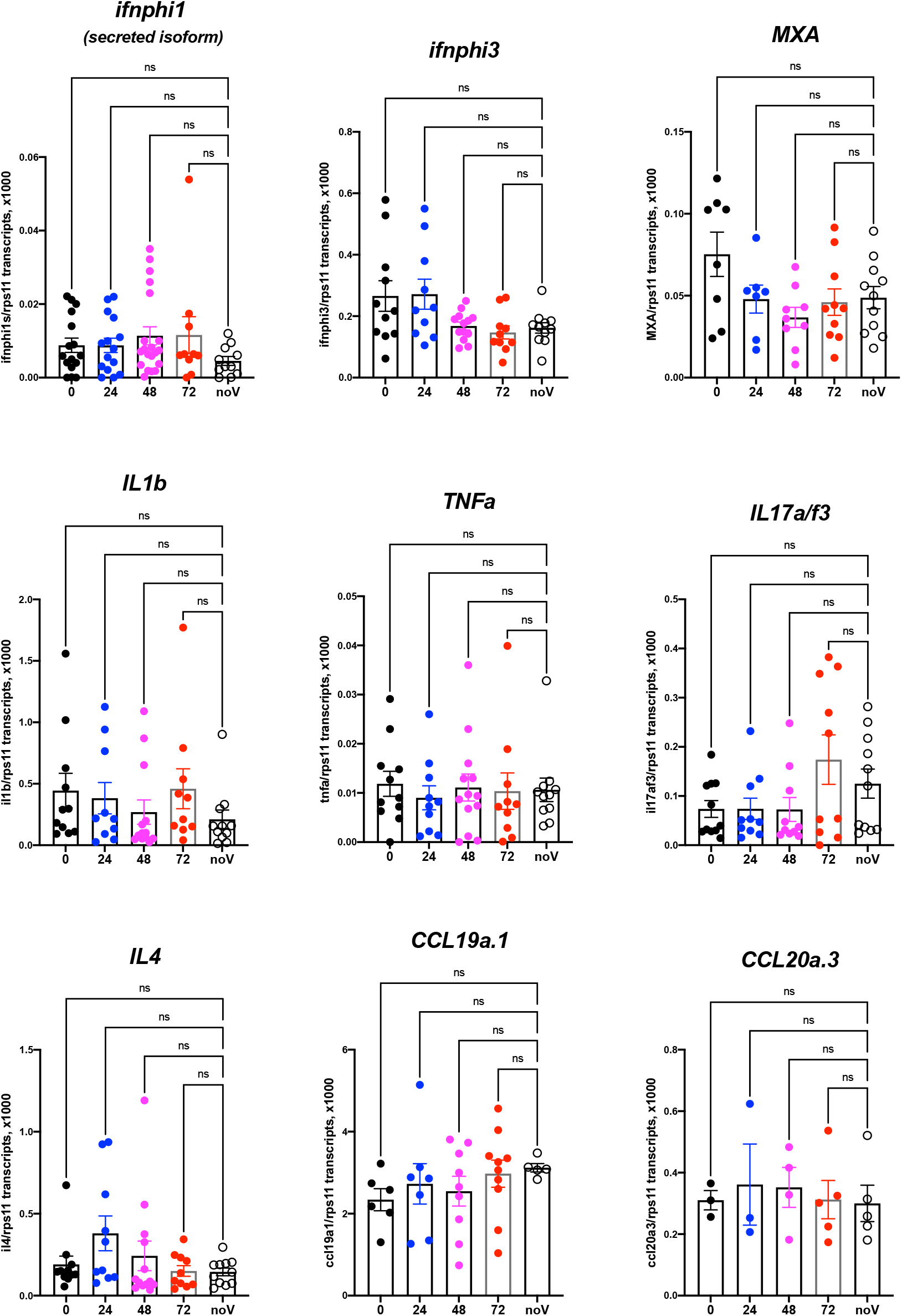
Host response after SARS-CoV-2 injection in the swim bladder. qRT-PCR, pool of 4 independent experiments (except for ccl19a.1and ccl20a.3, 3 and 2 experiments respectively). Numbers on X axis refers to hours post injection; noV (for “no Virus”): pooled uninjected negative controls, age-matched to 24, 48 or 72 hpi. One-way ANOVA analysis.

Although these results do not exclude a local response to SARS-CoV-2, they are in striking contrast with the those we obtained previously in larvae infected with other pathogens such as SINV or *Shigella flexneri*, for which many of these genes were induced more than 100-fold (Boucontet et al., 2018). Since these experiments had been performed at 28°C, we verified that zebrafish larvae are also able to mount a strong type I response at 32°C (Figure S3).

### Mosaic overexpression of hACE2 is not sufficient to support SARS-CoV-2 infection of 3 dpf larvae or fish cells in vitro

Finally, we tested if mosaic overexpression of human ACE2 in zebrafish larvae would increase their infectivity of SARS-CoV-2. We subcloned the *hace2* ORF in fusion with mCherryF under the control of the promoter of the ubiquitous ribosomal protein RPS26. In addition, the fragment is flanked by two inverted I-SceI meganuclease sites for higher transgenesis efficiency (Grabher et al., 2004). In order to be sure that the in-frame fusion of hACE2 with mCherry would not interfere with SARS-CoV2 binding to its receptor and entry in the target cells, another construct was done by inserting a self-cleaving 2A peptide between hACE2 and mCherry ORFs. We optimized the injected dose of plasmid; 68 pg was the amount yielding the highest mCherry expression without increasing the proportion of misshapen embryos (Figure S4A). In 24 hpf embryos, many mCherry^+^ cells, randomly distributed, were visible in these embryos under the fluorescence microscope. In swimming larvae, mCherry^+^ cells were still clearly visible but in lower amounts (Figure S4B). To get a quantitative assessment of their frequency, we dissociated 4dpf larvae and analyzed the suspension by flow cytometry, which indicated that ∼0.5% of the cells were mCherry^+^(Figure S4C). Larvae were fixed and processed by immunohistochemistry, which confirmed ACE2 expression at the membrane of mCherry^+^ cells (Figure S4D).

Zebrafish AB eggs were injected with the plasmid, and at 3dpf, the 25% larvae displaying the highest mCherry expression and good morphology were selected. They were then microinjected with SARS-CoV-2 in the coelom or the brain ventricle, and processed as above. qRT-PCR analysis revealed that viral mRNA transcripts decreased just as it did in AB larvae (Figure 9). Thus, this approach did not increase infectivity of SARS-CoV-2 in zebrafish larvae.

**Figure 9.**
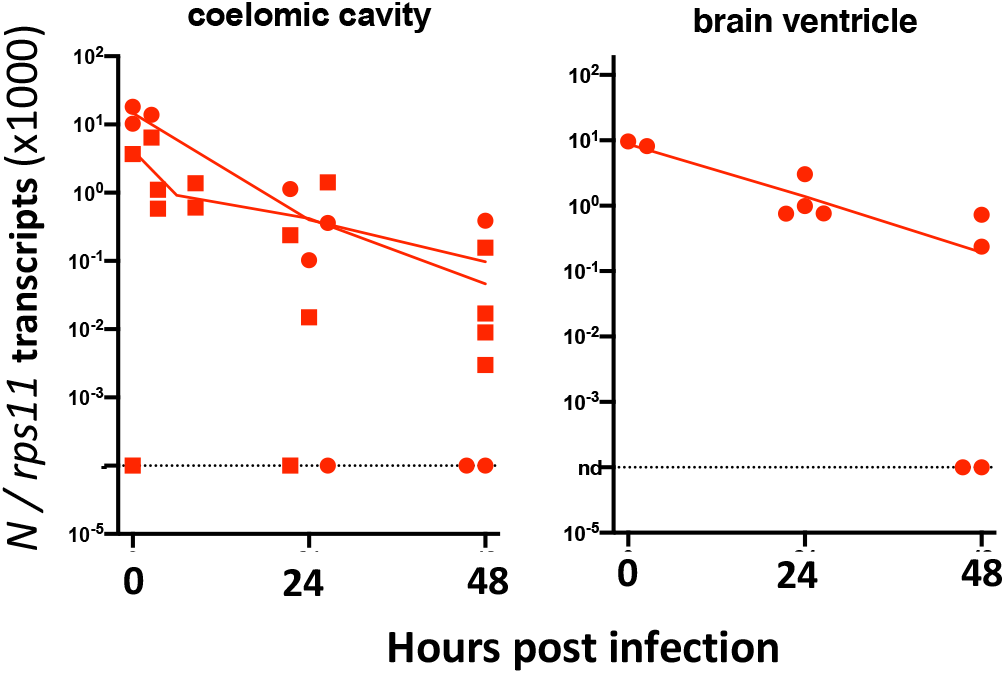
Injection of 3dpf hACE2-mCherry mosaic larvae. Quantification of sense N transcripts in individual hACE2-mCherry mosaic larvae injected in coelomic cavity (left; one experiment with hACE2-mCherry, one with hACE2-2A-mCherry) or brain ventricle (right; with hACE2-2A-mCherry) by qRT-PCR.

We finally tested if hACE2 overexpression by *in vitro* cultured fish cells made them susceptible to SARS-CoV-2, using the cyprinid cell line EPC. EPC cells were co-transfected with GFP and hACE2 expression plasmids; transfection efficiency and membrane hACE2 expression was verified by IHC (Figure S5A,B). These transfected cells were incubated with active or heat-killed SARS-CoV-2 at a MOI of 0.1, and then tested for viral replication by qRT-PCR on cell lysates. No difference was observed between GFP-only and GFP+hACE2 expressing cells (Figure S5C); furthermore, the amount of N transcripts fell dramatically from day 0 to day 2, showing that hACE2-expressing EPC cells were not able to support SARS-CoV-2 replication.

## Discussion

We report here our in-depth attempt to infect zebrafish larvae with SARS-CoV-2. Only larvae were tested because they present multiple practical advantages over adult fish: they can be rapidly generated in large quantities, incubated in multi-well plates, are highly amenable to imaging, and subject to fewer ethical regulations; therefore, they would be most suitable to drug screening. Whether juvenile or adult zebrafish would be more susceptible to SARS-CoV-2 remains to be tested.

We used absolute qRT-PCR of viral transcripts to test for viral replication. Surprising high numbers were measured shortly after injection, as the concentrated viral suspensions we used contained a considerable amount of non-infectious viral molecules, including negative strand species. In all likelihood, these molecules were released by infected Vero-E6 cells during the production of the virus stock; possibly by living cells as defective viral particles or in vesicles such as exosomes, or as free or membrane-bound RNA from dying cells. Whatever their origin, they complicate the detection of active viral replication, which has to generate enough molecules to exceed this background.

In almost all of our tests, a rapid (10 to 100-fold) decrease of mRNA copies was observed during the first day, likely due to degradation of non-infectious RNA species. After a few hours bath exposure, viral RNA was detected in doubly-rinsed larvae; this did not require active fusion or viral particles as RNA was also detected after exposure to heat-inactivated virus, and may have resulted from sticking of particles to skin surfaces or entry in the pharyngeal cavity. Two days after the starting of exposure, viral RNA was undetectable and thus the virus failed to achieve infection by bath, consistent with the results of (Kraus et al., 2020).

Microinjection is the most common way to infect zebrafish larvae with viruses (Levraud et al., 2014). After microinjection of a few nanoliters in larvae, the inoculum was readily measurable; however, when injected in the coelom, the pericardium, the bloodstream or the brain ventricle, viral RNA copy numbers then steadily declined, indicating unsuccessful infection. Two injection sites yielded different results: the yolk and the swim bladder. In the yolk, no RNA decrease was observed, suggesting that viral RNA molecules – perhaps owing to their coating with nucleoprotein and/or their localization in vesicles – were spared from degradation. Importantly, the yolk was unique among all tested sites as the one where injection is performed inside the cytosol of a cell (the yolk syncytial cell, not to be confused with the yolk sac) and not in the extracellular milieu. This does not necessarily prevent infection, as other viruses, such as CHIKV (Briolat et al., 2014) or human noroviruses (Van Dycke et al., 2019) have been shown to infect larvae after yolk injection. No signs of yolk infection (such as opacity observed with CHIKV and SINV) were observed, and no increase of viral mRNA was observed, so we believe that yolk injection did not result in active SARS-CoV-2 replication.

By contrast, injections in the swim bladder resulted in a ∼20-fold decrease of mRNA copies during the first day, followed by a small re-increase of the mean associated with a strong dispersion of values. This strongly suggests that successful infection occurred in some but not all larvae after swim bladder infection. Replication remained modest however, with only a 2- to 3-fold increase in copy numbers per day. Because of the considerable spread in measured copy numbers at 2 dpi, the re-increase is statistically borderline, but the bimodal distribution observed in the independent type I IFN mutant assay, and the comparisons with injections in the coelom, support this finding. Importantly, this was also confirmed by an independent immunohistochemistry assay as we observed, in a fraction of injected larvae, a few cells in the swim bladder wall there labelled by an antibody that detects the SARS-COV-2 nucleoprotein. It remains unclear why infections succeed in only a fraction of swim-bladder injected larvae. This could be due to a very low effective inoculum, but this seems unlikely since the success rate was not obviously higher with viral suspension 1 than suspension 2, despite a 7-fold higher titer.

It is interesting that the organ found to be most permissive to infection in zebrafish larvae is homologous to the human lung which is the primary target of the virus. We do not know if swim bladder epithelial cells express *ace2*. Unfortunately, there is no “swim bladder epithelium” subset in the scRNAseq zebrafish developmental atlas (Farnsworth et al., 2020), perhaps because these cells are too rare or difficult to isolate enzymatically. However, the swim bladder derives from the gut, which is the only organ in which cells highly express *ace2* in the atlas (Postlethwait et al., 2020). One may speculate that, besides surface protein expression, biophysical parameters such as surfactant coating or pressure-mediated tension of the epithelium could contribute to infectivity.

Not surprisingly, the SARS-COV-2 virus has evolved during the pandemic with successful waves of variants of concern with mutated spike protein, predicted to modulate binding to hACE2 and antibody neutralization. In the normally non-permissive wild-type mouse model, it has been shown that the beta and gamma variants replicated to a significant extent (Montagutelli et al., 2021). We tested several variants, including those two, in the zebrafish swim bladder model but did not find increased infectivity compared to the reference strain.

To stay within the thermal range of both virus and host, we incubated SARS-CoV-2-injected larvae at 32°C. Because SARS-CoV-2 replicates better at 33°C than 37°C in mammalian cells (V’kovski et al., 2020) (and our own observations), this is unlikely to be the reason for the poor replication of the virus in larvae. We also verified that at this temperature, larvae are able to mount a type I IFN response against another virus, eliminating temperature stress as the explanation for the lack of inflammatory response of zebrafish larvae to SARS-CoV-2. This is more likely a due to the small number of infected cells in our conditions, and possibly also active inhibition of some innate immune pathways by the virus. Protocols resulting in stronger infection will be needed for studying SARS-CoV2-induced inflammation in zebrafish larvae. This absence of measurable type I IFN response is consistent with the finding that IFN or IFN-R deficiency did not rescue virus infectivity. Thus, a limited compatibility between the virus and the host, rather than an intrinsic active resistance, seems the most likely explanation for our largely negative results.

Mosaic overexpression of hACE2 did not result in infectivity of 3 dpf larvae by SARS-CoV-2. We do not know if this was due to the relatively small number of cells expressing the transgene (<1%), to low expression or misfolding of the hACE2 protein, and/or to other causes. As an alternative strategy, we also tested injection of synthetic mRNA encoding hACE2-mCherryF; this resulted in clear ubiquitous mCherry expression at 24 hpf, but it had become undetectable by 2 dpf (not shown). This suggests that the hACE2 protein has a relatively short half-life in the zebrafish larval context. This issue may be solved by the establishment of stable transgenic zebrafish lines expressing hACE2. However, we also tested the effect of overexpression of hACE2 in the more stable context using the EPC cell line. EPCs are derived from a cyprinid fish, and used routinely to test the pro- of anti-viral activity of zebrafish genes by overexpression (e.g., (Langevin et al., 2013)). However, expression of hACE2 was not sufficient to allow replication of SARS-COV-2 on these cells. The lack of replication may be due to the need for co-expression of the transmembrane serine protease TMPRSS2, which has been shown to greatly increase SARS-CoV2 infectivity (Hoffmann et al., 2020). We also attempted to overexpress human TMPRSS2 in zebrafish embryos, either by plasmid or mRNA injection; unfortunately, this was found to be highly toxic, as it resulted in severe developmental anomalies that precluded injections.

In conclusion, our experiments indicate that the zebrafish larva is largely not infectable by SARS-CoV-2, except when the virus is injected in the swim bladder, which appears to result in modest viral replication in a subset of the animals. Given the expression of *ace2* in zebrafish enterocytes, it would also have been interesting to microinject the virus in the gut lumen. We tried, unsuccessfully, in part because of the close apposition of the gut and the easily damaged yolk. It should be noted however, that coelomic injections (the equivalent of intraperitoneal injections), comparatively easy to perform, deliver the virus in close proximity to the basal side of enterocytes, but do not yield successful infection. Further optimization of infection procedures, starting with the generation of transgenic zebrafish expression stably expressing human ACE2, will be needed to unleash the full potential of the zebrafish larva in the fight against COVID-19.

## Methods

### Ethical statement

Animal experiments described in the present study were conducted according to European Union guidelines for handling of laboratory animals (http://ec.europa.eu/environment/chemicals/lab_animals/home_en.htm) and were Approved by the Ethics Committee of Institut Pasteur.

### Fish

Wild-type zebrafish (AB strain), initially obtained from ZIRC (Oregon, USA) were raised in the aquatic facility of Institut Pasteur. After natural spawning, eggs were collected, treated for 5 minutes with 0.03% bleach, rinsed twice, and incubated at 28°C in Petri dishes in Volvic mineral water supplemented with 0.3μg/mL methylene blue (Sigma-Aldrich). After 24 hours, the water was supplemented with 200μM phenylthiourea (PTU, Sigma-Aldrich) to prevent pigmentation of larvae. After this step, incubation was conducted at 24, 28 or 32°C depending on the desired developmental speed. Developmental stages given in the text correspond to the 28.5°C reference (Kimmel et al., 1995). Sex of larvae is not yet determined at the time of experiments.

Triple type I interferon CRISPR mutants have been generated by the AMAGEN transgenesis platform (Gif-sur-Yvette, France) by co-injection of CAS9 with two sgRNA targeting *ifnphi1* (target sequence, GCTCTGCGTCTACTTGCGAAtgg) and *ifnphi2* (target sequence, ATGTGCGCGAAAAAGAGTGCtgg) in one-cell eggs from homozygous ifnphi3^ip7/ip7^ null mutants of AB background (Maarifi et al., 2019). After growth to adulthood, a founder was identified that co-transmitted mutations in *ifnphi1* and *ifnphi2* in addition to the *ip7* mutation of *ifnphi3*. The *ip9* allele mutation in *ifnphi1* consists in a 7 base pair deletion in the first exon of the secreted isoform (GAATGGC, 75 bases downstream of the start ATG). The *ip10* allele in *ifnphi2* consists in a 19bp deletion in the first exon (TGCGTTCTTATGTCCAGCA, 20 bases downstream of the start ATG). This founder was crossed with AB fish and F1 fish triply heterozygous for mutations *ip7, ip9* and *ip10* were selected to establish the “triple φ” mutant line. As expected since *ifnphi1, ifnphi2* and *ifnphi3* are closely located in tandem on a 35 kb region of zebrafish chromosome 3, the *ip7, ip9 and ip10* mutations were always found to co-segregate. Genotyping PCR primers are listed on Table 2.

**Table 2.**
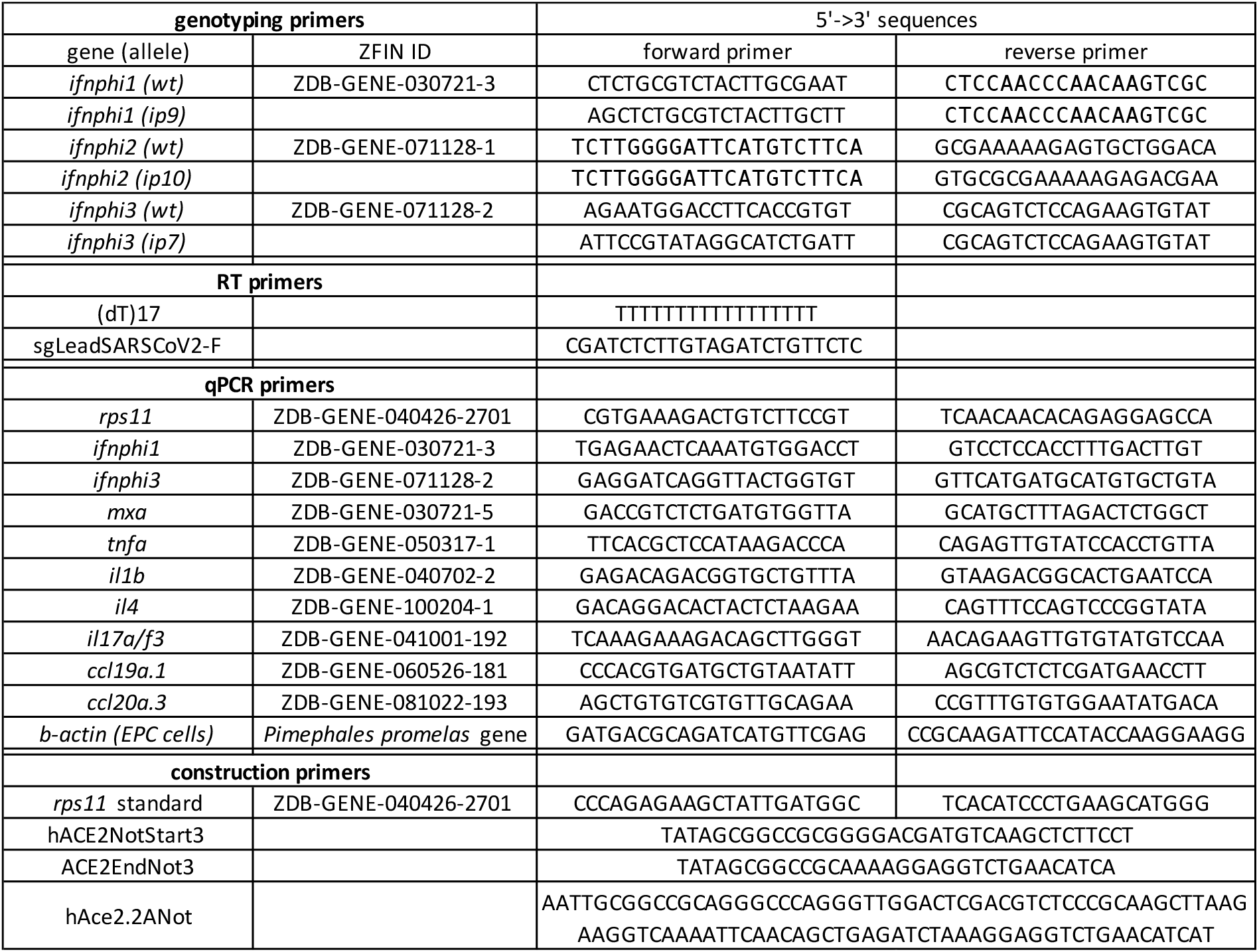
Primers used in this study.

### Viruses

The main SARS-CoV-2 stock used (BetaCoV/France/IDF0372/2020 strain) was propagated twice in Vero-E6 cells and is a kind gift from the National Reference Centre for Respiratory Viruses at Institut Pasteur, Paris, headed by Dr Sylvie van der Werf; this strain was isolated from a human sample provided by Drs. Xavier Lescure and Yazdan Yazdanpanah from the Bichat Hospital, Paris. To generate concentrated virus, Vero-E6 cells were infected with virus at an MOI of 0.01 PFU/cell in DMEM/2%FBS, and incubated for 72 h at 37°C, 5% CO2. At this point, the cell culture supernatant was harvested, clarified and concentrated using Amicon Ultra-15 Centrifugal units 30K (Merck Millipore). Virus titers were quantified by plaque assay in Vero-E6.

The variants strains used were also supplied by the National Reference Centre for Respiratory Viruses at Institut Pasteur and were used directly without further propagation. The G-clade (BetaCoV/France/GE1973/2020; 3×10^7^ PFU/ml), alpha (hCoV-19/France/IDF-IPP11324i/2020; 6.75×10^7^ PFU/ml), beta (hCoV-19/France/PDL-IPP01065i/2021; 1.75×10^8^ PFU/ml), and gamma (hCoV-19/French Guiana/IPP03772i/2021; 5.53×10^7^ PFU/ml) variants were isolated from human samples provided respectively by Dr Laurent Andreoletti, from Robert Debré Hospital, Reims, France; Dr Foissaud, HIA Percy, France; Dr Besson J. from Bioliance Laboratory, France; and Dr Rousset, Institut Pasteur, Cayenne, French Guiana.

The SINV-GFP virus corresponds to the SINV-eGFP/2A strain described in (Boucontet et al., 2018) and was used as a BHK cell supernatant at 2×10^7^ PFU/mL.

### Bath exposure

Bath exposures were conducted in a 12-well plate with 4 larvae per well in 2 mL of water plus PTU. 2dpf embryos were manually dechorionated previously. 10 μL of SARS-CoV-2 suspension 2 (either freshly thawed or heat-inactivated for 5 minutes at 70°C) was added to each well and gently mixed, then the plates were incubated at 32°C. After a given incubation time, larvae were deeply anesthetized with 0.4mg/ml tricaine (MS222, Sigma-Aldrich), rinsed twice in 10mL of water, transferred individually into tubes and after removal of almost all water, lysed in 320 μl of RLT buffer (Qiagen) supplemented with 1% β-mercaptoethanol (Sigma-Aldrich).

### Microinjection

SARS-CoV-2 microinjections are carried out under a microbiological safety hood inside a BSL3 laboratory, in which a camera-fitted macroscope (DMS1000, Leica) with a transilluminated base is installed, as in (Van Dycke et al., 2019). Borosilicate glass capillaries are loaded with a concentrated SARS-CoV-2 suspension previously coloured by the addition of 10% (V/V) of 0.5% phenol red in PBS (Sigma), then connected to a FemtoJet 4i microinjector (Eppendorf). Otherwise, the procedure was similar to the one detailed in (Levraud et al., 2008). After breakage of the capillary tip, pressure was adjusted to obtain droplets with a diameter of ∼0.13mm. Larvae at the desired developmental stage were anesthetized with 0.2mg/mL tricaine and positioned and oriented in the groove molded in agarose of an injection plate overlaid with water containing tricaine. Using a micromanipulator, the capillary was then inserted at the desired site and two pulses performed to inject approximatively 2 nL. Proper injection is ascertained visually with the help of phenol red staining; otherwise, the larva is discarded. A picture of the injected larva is taken with the camera, and it is then rinsed by transfer inside a water-filled Petri dish and immediately transferred to its individual well in a 24-well plate, containing 1mL of water with PTU. Larvae are then incubated at 32°C (actual temperatures measured inside the incubator ranged from 31.6 to 33.2°C). At daily intervals, all larvae were anesthetized by addition of a drop of 4mg/mL tricaine into each well and a snapshot was taken. A randomized subset of larvae was then transferred to tubes and individually lysed in 320 μl of RLT buffer + 1% β-mercaptoethanol. Water with tricaine was then removed from the remaining wells, replaced with 1 mL of fresh water with PTU, and the plate returned to the incubator.

SINV injections were performed in a BSL2 laboratory as described in (Passoni et al., 2017).

### Lysis, RNA extraction, and RTqPCR of larvae

After addition of RLT buffer, larvae were dissociated by 5 up-and-down pipetting movements. Tubes may then be frozen at -80°C for a few days. Before export from the BSL3 laboratory, RLT lysates were incubated at 70°C for 5 minutes to ensure complete virus inactivation (preliminary tests confirmed that this had a negligible impact on qRT-PCR results). Total RNA was then extracted with a RNeasy mini kit (Qiagen) without the DNAse treatment step and a final elution with 30μL of water.

RT was performed on 6μl of eluted RNA using MMLV reverse transcriptase (Invitrogen) with either a dT_17_ primer (for polyadenylated transcripts) or the SgleadSARSCoV2-F primer (for negative strand viral transcripts) (Wölfel et al., 2020)(Table 2). cDNA was diluted with water to a final volume of 100 μL, of which 5 μL was used as a template for each qPCR assay.

Real time qPCR was performed with an ABI7300 (Applied Biosystems). Quantitation of sense or antisense viral *N* transcripts was performed by a Taqman probe assay, using the primer-probe mix from the 2019-nCoV RUO kit (IDT) with iTaq Universal Probes One-Step kit (Bio-Rad). The 2019-nCoV_N_Positive Control plasmid (IDT) was used as a standard for absolute quantification. Quantification of zebrafish transcripts was performed using a SYBR assay using the Takyon SYBR Blue mastermix (Eurogentec) with primer pairs listed on Table 2. These primers typically span exon boundaries to avoid amplification of contaminating genomic DNA. For absolute quantification of the housekeeping gene *rps11*, a standard was produced by PCR using primers to amplify a fragment including the whole open-reading frame, which was gel-purified and quantified by spectrophotometry. Ratios of other transcripts to *rps11* were estimated by the 2^ΔCt^ method.

### Morpholino and plasmid injection in eggs

Morpholino antisense oligonucleotides (Gene Tools) were injected (1 nL volume) in the cell or yolk of AB embryos at the one to two cells stage as described (Levraud et al., 2008). crfb1 splice morpholino (2 ng, CGCCAAGATCATACCTGTAAAGTAA) was injected together with crfb2 splice morpholino (2 ng, CTATGAA TCCTCACCTAGGGTAAAC), knocking down all type I IFN receptors (Aggad et al., 2009). Control morphants were injected with 4 ng control morpholino, with no known target (GAAAGCATGGCATCTG GATCATCGA).

Expression plasmids, produced using an endotoxin-free kit (Macherey-Nagel), were co-injected with the I-SceI meganuclease (Grabher et al., 2004). Briefly, 12.5μL of plasmid is mixed with 1.5μl of Custmart buffer and 1μl of I-SceI (New England Biolabs), and incubated at room temperature for 5 min before being put on ice until injection of 1 nL inside the cell of AB embryos at the one-cell stage.

### Live fluorescence imaging

SINV-GFP infected or hACE2-mCherry expressing larvae were imaged with an EVOS FL Auto microscope (Thermo Fisher Scientific) using a 2× planachromatic objective (numerical aperture, 0.06), allowing capture of the entire larva in a field. Transmitted light and fluorescence (GFP or Texas Red cube) images were taken. They were further processed (superposition of channels, rotation, crop, and fluorescence intensity measure) using Fiji. Mean background fluorescence of uninjected control animals was subtracted from the measured signal to obtain the specific fluorescence.

### Flow cytometry

Pools of 10 larvae were dissociated by a combination of mechanical trituration (repeated pipetting) and enzymatic treatment at 30°C, first with 200μL of 0.25% Trypsin-EDTA (Gibco) for 10 minutes, and 10 more minutes after addition of 10% sheep serum, CaCl2 to 2μM, and 1μL of 5mM collagenase (C9891, Sigma). Cell suspensions were then washed with PBS 1x, pelleted, resuspended in PBS, and filtered on a 40μm mesh. Dead cells were labelled with Sytox AADvanced (ThermoFisher). Cell suspensions were acquired on an Attune NxT flow cytometer (ThermoFisher) with blue and yellow lasers, and data analyzed with FlowJo.

### Cell culture

*Epithelioma papulosum cyprini* cell line (EPC) was maintained in Leibovitz-15 media (L15, Gibco) supplemented with 10% fetal bovine serum (FBS, Gibco), 100 μg/l penicillin and 100 μg/ml streptomycin. EPC cells were cultured at 32 °C without CO_2_.

### Human ACE2 expressing constructs

The hACE2 ORF was amplified from clone IOH80645 (Thermosfisher, GenBank NM_021804.2) using primers hACE2NotStart3 and hACE2EndNot3 (table 2). The amplified PCR fragment was digested by NotI and inserted in the NotI site of the Tol2S263C:mC-F vector between the promoter of the zebrafish ubiquitous ribosomal protein RPS26 encoded by chromosome3, and the mCherry ORF. In this construct, the RPS26 promoter drives the expression of a hACE2 protein fused at its C-term with farnesylated mCherry. In order for the ORF to drive the expression of two separated proteins (hACE2 and mCherry-F), primers hACE2NotStart3 and hAce2.2ANot were used to amplify the hACE2 ORF from the IOH80645 clone. The amplified fragment was digested by NotI and cloned in the NotI site of Tol2S263C:mC-F between the promoter of the zebrafish ubiquitous ribosomal protein RPS26 encoded by chromosome3, and the mCherry ORF. Maps and sequences of plasmids are available at https://doi.org/10.5281/zenodo.4672028.

For *in vitro* transfection of EPC cells, plasmid pcDNA3.1-hACE2 (Addgene #1786) was directly used along plasmid pmEGFP-N1 (Chen & Reich, 2010).

### Cell transfection

EPC cells were electroporated with the Neon transfection system (Invitrogen). Briefly, EPC cells were trypsinized and resuspended in L15 media supplemented with FBS and antibiotics. Cells were counted and centrifugated at 2000 rpm for 5 minutes. 0.8 × 10^6^ cells per condition were resuspended in 80 μl of L15 without phenol red with 2.4 μg of each plasmid. Cells were electroporated using 10 μl neon tips with 1 pulse of 1700 V during 20 ms. Electroporated cells were plated in a 6-well plate in L15 + FBS + antibiotics and incubated 3 days at 32°C before experiment.

### Cell infection and RT-qPCR

Transfected EPC cells were transferred to BSL3 laboratory for infection with SARS-CoV2. Cells were rinsed with L15 media + FBS + antibiotics and incubated 5 minutes at 32°C. Cells were infected at MOI 0.1 with virus diluted in L15 media + FBS + antibiotics and incubated at 32°C during 1 hour with agitation. After incubation, L15 media + 10% FBS + antibiotics was added and cells were incubated 2 days at 32°C or processed directly for RNA extraction.

Before RNA extraction, culture medium was removed and cell were rinsed once with PBS. Extraction of total RNA was performed using Tri-Reagent (Sigma) following manufacturer recommendations. Total RNA was resuspended in 100 μl of RNase-free water.

Reverse transcription was performed on 5 μl of RNA suspension using QuantiTect Reverse Transcription kit (Qiagen) with either the qiagen RT primer mix or the SgleadSARSCoV2-F primer (for negative strand viral transcripts) (Wölfel et al., 2020). cDNA was diluted with water to a final volume of 50 μL, of which 2.5 μL was used as a template for each qPCR assay.

Real time qPCR was performed with a Realplex2 (Eppendorf). Quantitation of viral N transcripts was performed by a Taqman probe assay, using the primer-probe mix from the 2019-nCoV RUO kit (IDT) with iTaq Universal Probes kit (Bio-Rad). Quantitation of actin transcripts was performed by a SYBR green assay, using primers specific for fathead minnow β-actin (Table 2) with iTaq universal SYBR green supermix (Bio-Rad).

### Immunohistochemistry

Whole-mount immunohistochemistry of larvae was performed essentially as described in (Palha et al., 2013) and (Santos et al., 2018). For COV2-N detection, additional treatment with glycine 0.3M in PBST (30 minutes at RT) and Heat induced antigen retrival (HIER) were performed. HIER treatment was performed in 150mM Tris-HCl, Ph 9.0 at 70 C for 15 min. Primary Ab used for this labelling were: mouse anti-SARS COV2 nuceloprotein (Sino Biological, 40143-MM05, 1:100) and rabbit anti-GFAP (GeneTex, GTX128741, 1:100). As secondary Ab were used: goat anti-mouse F(ab)’2 AlexaFluor 488 (Molecular Probes, A11017, 1:300) and goat anti-rabbit Cy3 (Jackson Laboratories 111-166-003, 1:300). Furthermore, to label the nuclei was used NucRed Live 647 (ThermoFisher, R37106, 4 drops for mL for 45 minutes). For hACE2 detection, stainings were performed sequentially since both the primary Ab for ACE2 and the secondary Ab for mCherry were from goat. Primary staining for ACE2 (goat anti-ACE2, AF933, R&D systems, 4μg/mL) was performed first, followed by its secondary staining (donkey anti-goat Ig Alexa 488, A100555, Invitrogen, 1:300), then primary staining for mCherry (rabbit anti-DsRed, 632393, Clontech, 1:300) and secondary staining (goat anti-rabbit Ig Cy3, 111-166-003, 1:300). Nuclei were labelled with 2μg/mL Hoeschst 33342 (Invitrogen).

After IHC larvae were conserved in 80% Glycerol until acquisition. For acquisition of N-CoV2 the larvae were mounted in 2% Agarose in 80% Glycerol singularly in a glass bottom 8 wells slide (Ibidi, 80827).Images were acquired using inverted confocal microscope Leica SP8 using 10x objective zoomed 1.25x (PL FLUOTAR 10x/0.30 DRY) and 20x immersion objective (HC PL APO CS2 20x/0.75 multi-IMM). For both magnification bidirectional resonant scanning method was used and images were deconvolved using Leica Lightening Plug-in. For acquisition of hACE2 images were acquired on an upright Leica SPE confocal microscope using a 40x oil objective (numerical aperture, 1.15).

For IHC of *in vitro* transfected cells, EPC were cultured in 6-well plate containing sterilized coverslips. At 3 days-post transfection, culture media was removed and cells were rinsed with PBS once. Cells were fixed overnight at 4°C with 4% methanol-free formaldehyde (Sigma) in PBS. Formaldehyde was removed and cells were rinsed twice with PBS and kept at 4°C in PBS + 0.05% sodium azide. Fixed cells were rinsed 3 times in PBS. Cells were then permeabilized and blocked with PBS + 0.3 % triton X-100 + 10% horse serum during 45 min at RT. Cells were stained 1h at RT with a goat polyclonal anti-human ACE2 (AF933, R&D Systems) diluted at 3μg/ml in PBS + 0.3% triton X-100 + 1% horse serum + 1% BSA + 0,01% sodium azide. Cells were then rinsed and stained during 1h at RT with Alexa647 anti-goat diluted at 1/500 in PBS + 0.3% triton X-100 + 1% horse serum + 1% BSA + 0.01% sodium azide. After 3 rinsing with PBS, cells were incubated 1h at RT with DAPI diluted at 2.5 μg/ml in PBS. After 3 rinses in PBS, coverslips were mounted on slides with Fluoromount G (ThermoFisher Scientific).

Transfection efficiency was checked at 3 days post-transfection using a Zeiss Axio Observer Z1 widefield microscope with a 10X/NA 0.25 objective. Phase and GFP channel were acquired on 5 field of view. Confocal acquisition of immunostained EPC cells was performed on a Leica SP8 upright microscope using a 25X/NA 0.95 coverslip corrected objective. Endogenous GFP and Alexa 647 were excited with 488 nm and 638 nm respectively and detected with PMT. Fiji was used to adjust brightness and contrast of confocal images of immunostained EPC cells. Transfection efficiency was quantified using Fiji by manually counting total cells and GFP expressing cells, respectively.

### Statistical analysis

Analysis were performed with GraphPad Prism. Methods used are indicated in Figure legends. Normality/lognormality tests of data distribution were performed to decide the most appropriate assays.

## Acknowledgements

We thank Frédéric Sohm and Joanne Edouard (AMAGEN, Gif-sur-Yvette) for the generation of the triple interferon mutants; our animal facility manager Yohann Rolin for ensuring fish health and fertility despite lockdowns; numerous members of Institut Pasteur for sharing materials (Olivier Schwartz, Julian Buchrieser, Pierre Charneau, Viviana Scoca, Amandine Noirat, Nathalie Sauvonnet, Pierre lafaye, Christophe Zimmer, Christian Weber), managing the BSL3 lab (Ferdinand Roesch, Florence Guivel-Benhassine, Thomas Vallet, Heloise Mary), occasional help in experiments (François Huetz, Björn Meyer, Doris-Lou Demy) and useful discussions (Emma Colucci-Guyon, Anne Schmidt, Philippe Herbomel). We thank Bertrand Collet and Lise Chaumont (INRAE, Jouy en Josas) for sharing the pmEGFP-N1 plasmid, helpful advice on transfection, and fathead minnow b-actin primers.

## Funding

This work was supported by the “Urgence COVID-19” fundraising campaign of Institut Pasteur, a dedicated grant co-funded by Institut Pasteur and Institut du Cerveau -Paris Brain Institute, the “ZebraCorona” grant from the exceptional research program call of Université Paris-Saclay, the MUSE-University of Montpellier COVID program, the Agence Nationale de la Recherche (Grant ANR-16-CE20-0002-03 « fish-RNAvax ») and the “ImageInLife” Innovative Training Network funded by European Community’s Horizon 2020 Marie-Curie Program under grant agreement n°721537.

## Competing interests

The authors declare no competing interests.

## Author contributions

JPL, PB, IS and MV designed the study, which was coordinated by JPL. VR generated the concentrated SARS-CoV-2 virus and supervised early BSL3 work in IP. BdC supervised BSL3 work in INRAE. JPL and VL performed 1-cell injections and SARS-CoV-2 microinjections in the BSL3 lab, VL performed SINV injections, WIHC, and fluorescence imaging. MF performed in vitro work. JPL, LB, VL and MF performed qRT-PCR assays. GL generated the overexpression plasmids. JPL wrote the manuscript with input from all authors.

## Notes

https://doi.org/10.5281/zenodo.4672028

### Competing Interest Statement

The authors have declared no competing interest.

### Summary of Updates

Confirmation of presence of infected cells in the swim bladder by whole-mount immunohostochemistry (figure 5, movie S1) Test of several variants of SARS-CoV-2 (Figure 6) Checking if overexpression of hACE2 allowed infection of a fish cell line in vitro (Figure S5)

