## supplemental Figures S1-S5 for "Exploring zebrafish larvae as a COVID-19 model: probable SARS-COV-2 replication in the swim bladder"

Supplementary figures S1-S5, with legends
Legend to movie S1

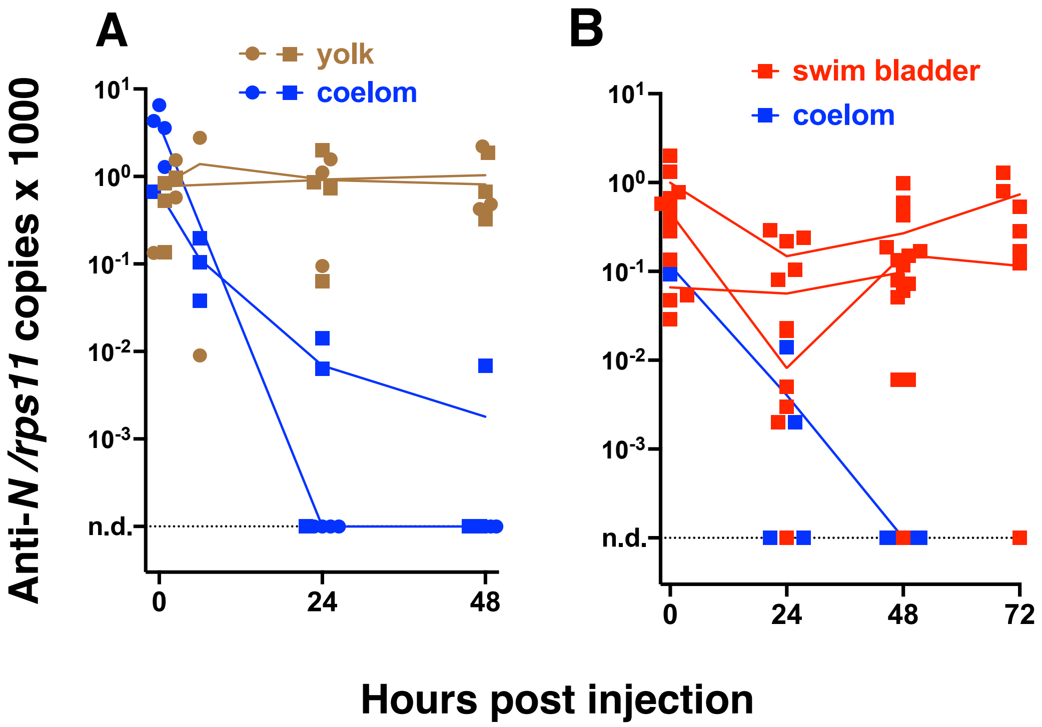


***Figure S1. Antisense viral transcripts in injected larvae**. qRT-PCR quantification of N viral transcripts with the leader sequence Each point corresponds to an individual larva. Lines connect means in each independent experiment. Circles and squares correspond to injection of viral suspensions 1 and 2, as labelled on Table 1, respectively. n.d., not detected. A. 3 dpf larvae injected in the coelom (blue) or yolk (brown) B. 4 dpf larvae injected in the coelom (blue) or swim bladder (red).*

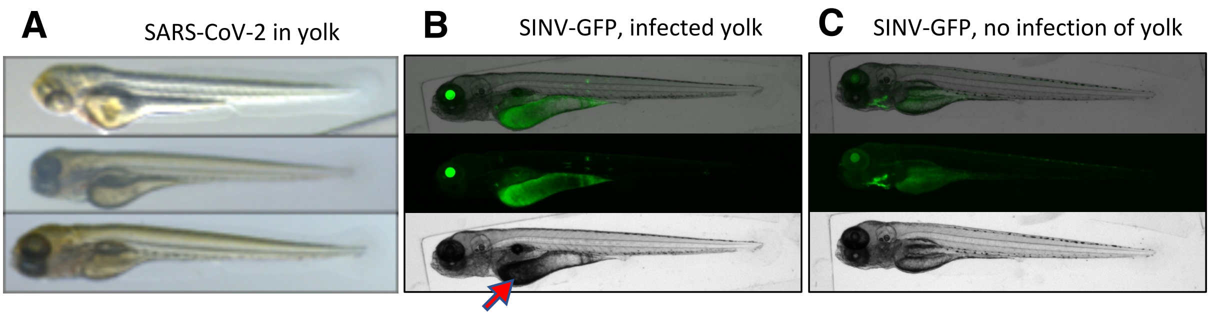


***Figure S2. Aspect of yolk-injected larvae.*** *A. transmitted light images of an individual larva injected with SARS-CoV-2 in the yolk just after injection (top), 24 (middle) and 48 hpi (bottom). B and C representative examples of SINV-infected larvae, 2 days after injection IV (B) and in the pericardium (C). GFP signal (middle), transmitted light image (bottom) and merged images (top). The GFP signal reveals localization of infection; red arrow points to the typical opacity of infected yolk.*

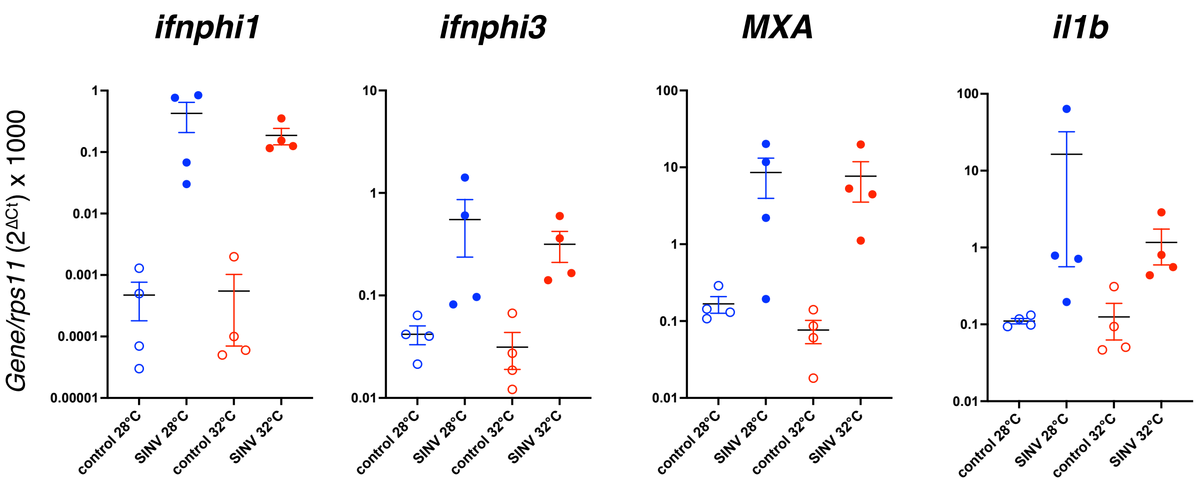


***Figure S3. Antiviral responses are inducible at 32°C.*** *qRT-PCR analysis of zebrafish larvae after 24 hours of incubation at either 28 (blue) or 32°C (red) following injection with 40 PFU of SINV-GFP. All pairwise comparisons (control vs SINV) were statistically different (p<0.05, Mann-Whitney test) except for MXA at 28°C (p=0.057)*

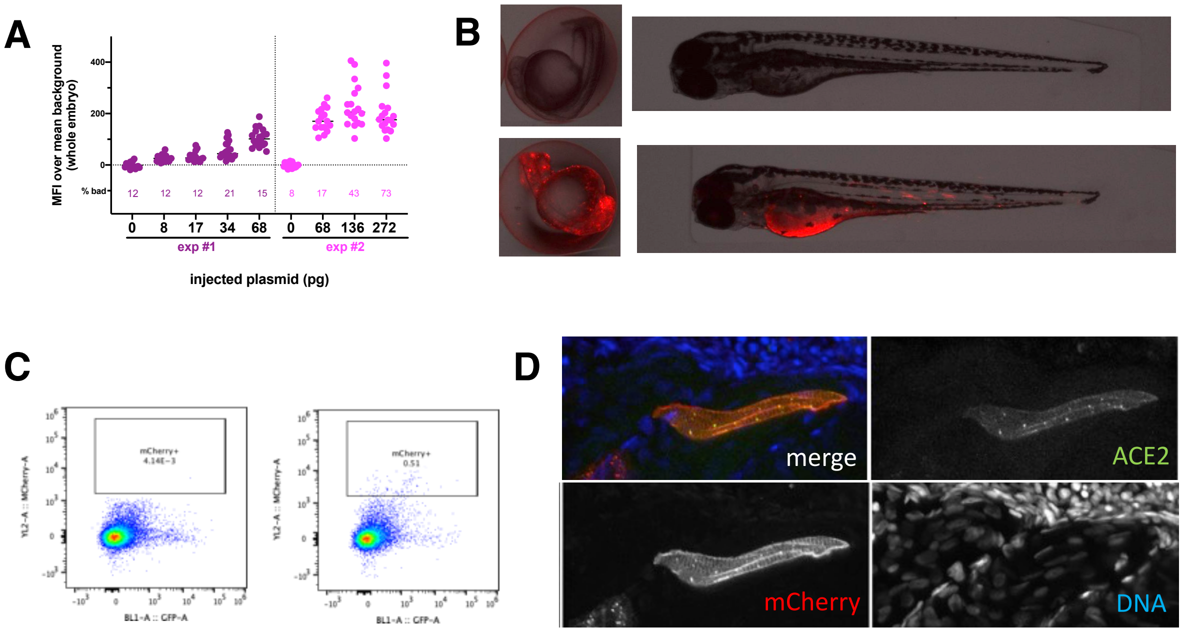


***Figure S4. Overexpression of hACE2-mCherry by plasmid injection at the 1-cell stage.*** *A. Optimization of the plasmid dose. Fluorescence intensity measured in 24hpf embryos after injection at the 1-cell stage of the specified amount of pz26hACE2-mCherryF plasmid together I-SceI, for the 25% embryos with best expression in each group. The percentage of misshapen embryos in each group is indicated on the bottom of the graph. B. representative image of a 24 hpf embryo (left) and of a 3 dpf larva (right) after mock injection (top) or injection of 68pg of pz26hACE2-mCherryF (bottom). C. Representative flow cytometry analysis of cells dissociated from 4dpf larvae, mock-injected (left) or injected with 68pg of pz26hACE2-mCherryF (right). D. Immunohistochemistry of a larva injected with 68pg of pz26hACE2-2A-mCherryF, showing ACE2 detection on a mCherry$^+$ muscle fiber.*

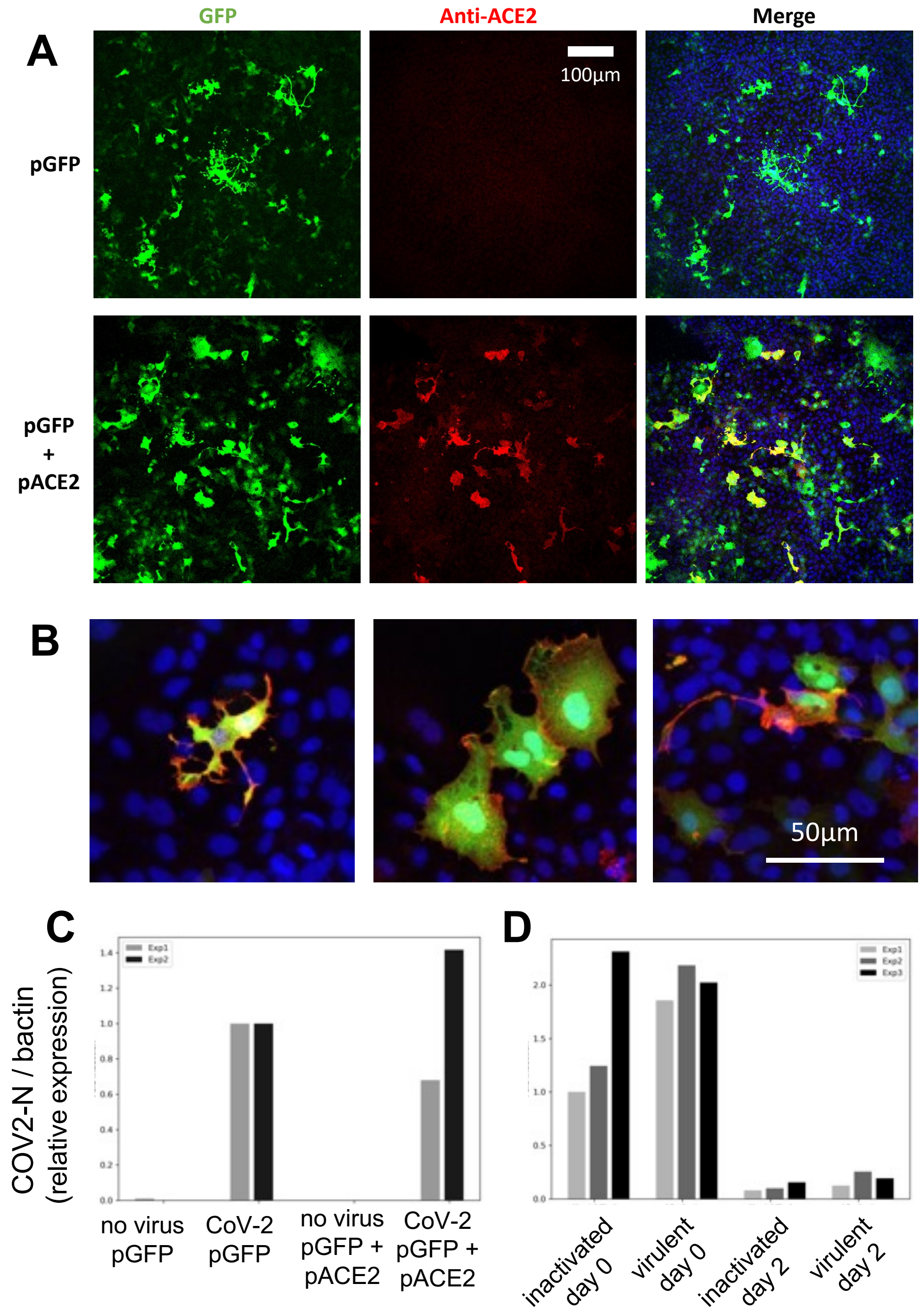


***Figure S5. SARS-CoV-2 does not replicate on EPC cells transfected with human ACE2.*** *A,B confocal microscopic images of EPC cells cultures 3 days after transfection. A. Assessment of transfection efficiency at low magnification; B. demonstration of hACE2 expression at the membrane of cells transfected with both plasmids, high magnification, merge images of GFP, hACE2 and nuclei labels. C, D qRT-PCR measurement of N copies in cells. C, measurement at 2 days post-exposure with virulent SARS-COV-2, comparison of GFP and GFP+hACE2 overexpression. D, decline of RNA levels from day 0 to day 2 post-exposure, comparison of heat-inactivated and virulent virus.*

**Movie S1. 3D reconstruction of a larva inoculated with SARS-CoV-2 in the swim bladder after immunodetection of CoV-2 nucleoprotein (green) and GFAP (red).** Nuclei are shown in blue. Related to Figure 5B.
